# Altered motility in response to iron-limitation is regulated by *lpdA* in uropathogenic *E. coli* CFT073

**DOI:** 10.1101/2023.09.27.559868

**Authors:** A.E. Frick-Cheng, A.E. Shea, J.R. Roberts, S.N. Smith, M.D. Ohi, H.L.T. Mobley

**Affiliations:** Life Sciences Institute, University of Michigan, Ann Arbor, MI, USA; Department of Microbiology and Immunology, University of South Alabama Medical School, Mobile, AL, USA; Department of Anesthesiology, University of Michigan Medical School, Ann Arbor, MI, USA; Department of Microbiology and Immunology, University of Michigan Medical School, Ann Arbor, MI, USA

**Author notes:** Address correspondence to Harry L. T. Mobley. equal contributions (A.E. Frick-Cheng, A.E. Shea).

## Abstract

More than half of all women will experience a urinary tract infection (UTI) in their lifetime with most cases caused by uropathogenic *Escherichia coli* (UPEC). Bacterial motility enhances UPEC pathogenicity, resulting in more severe disease outcomes including kidney infection. Surprisingly, the connection between motility and iron limitation is mostly unexplored, despite the lack of free iron available in the host. Therefore, we sought to explore the potential connection between iron restriction and regulation of motility in UPEC. We cultured *E. coli* CFT073, a prototypical UPEC strain, in media containing an iron chelator. Under iron limitation, CFT073 had elevated *fliC* (flagella) promoter activity, driving motility on the leading edge of the colony. Furthermore, this iron-specific response was repressed by the addition of exogenous iron. We confirmed increased flagella expression in CFT073 by measuring *fliC* transcript, FliC protein, and surface-expressed flagella under iron-limited conditions. To define the regulatory mechanism, we constructed single knockouts of eight master regulators. The iron-regulated response was lost in *crp, arcA,* and *fis* mutants. Thus, we focused on the five genes regulated by all three transcription factors. Of the five genes knocked out, the iron-regulated motility response was most strongly dysregulated in an *lpdA* mutant, which also resulted in significantly lowered fitness in the murine model of ascending UTI. Collectively, we demonstrated that iron-mediated motility in CFT073 is regulated by *lpdA*, which contributes to the understanding of how uropathogens differentially regulate motility mechanisms in the iron-restricted host.

**Importance:** Urinary tract infections (UTIs) are ubiquitous and responsible for over five billion dollars in associated health care costs annually. Both iron acquisition and motility are highly studied virulence factors associated with uropathogenic *E. coli* (UPEC), the main causative agent of uncomplicated UTI. This work is innovative by providing mechanistic insight into the synergistic relationship between these two critical virulence properties. Here, we demonstrate that iron limitation has pleiotropic effects with consequences that extend beyond metabolism, and impact other virulence mechanisms. Indeed, targeting iron acquisition as a therapy may lead to an undesirable enhancement of UPEC pathogenesis through increased motility. It is vital to understand the full breadth of UPEC pathogenesis to adequately respond to this common infection, especially with the increase of antibiotic resistant pathogens.

## Introduction

Urinary tract infections (UTIs) are a prominent health concern worldwide; they are the second most frequent infectious disease and disproportionately affect women, children, the elderly, immune-compromised, and hospitalized populations (1–3). Most uncomplicated UTIs are caused by uropathogenic *Escherichia coli* (UPEC) and are restricted to the bladder (4, 5). However, if left untreated, these infections can ascend the ureters to the kidneys causing permanent tissue damage and renal scarring; 80% of cases of pyelonephritis are caused by UPEC (6–8). In extreme cases, bacteria can further spread into the bloodstream potentially progressing to sepsis (8, 9). Indeed, about 25% of sepsis cases originate from UTIs (10). Even in the less severe, non-ascending cases of UTI, the financial burden and decreased quality of life are damaging to these patients (3, 11).

One of the most essential and well-studied mechanisms of bacterial dissemination from the bladder to the kidneys is motility (1, 12). UPEC has been detected in murine kidneys as early as 6 hours post-inoculation into the bladder (12). Flagella, expressed on the bacterial surface, propel the bacteria in a targeted direction utilizing chemotactic mechanisms (13, 14). Many studies have been conducted in prototype commensal strains of *E. coli* to understand the mechanical function and regulation of flagella and the chemoattractant stimuli that facilitate directed movement (15, 16). However, given that these studies have largely used non-pathogenic strains of *E. coli*, the established dogma of flagellar movement and regulation may differ in pathogenic varieties.

Strains of *E. coli* that cause human disease are distinctly genetically diverse from non-pathogenic serotypes (17, 18). For example, Trg and Tap chemoreceptors, which direct bacteria towards ribose, galactose and dipeptides (19, 20), are less prevalent and functional among UPEC isolates than fecal or commensal strains, and even more so when compared to diarrheal strains of *E. coli* (21). Because the genetic and phenotypic display of these pathogens are varied, it is likely that regulatory mechanisms of key processes that include bacterial motility, are too. Another example of divergence between pathogenic and commensal *E. coli* strains is the prevalence of iron importation systems. Uropathogens can import iron using multiple siderophore systems not found commensal isolates (22, 23). Iron acquisition is especially important for pathogens as the human host is an iron-poor environment (24). The ferric uptake regulator, Fur, was found to have additional binding sites within the pathogenicity islands of UPEC compared to commensal strains, including many of these additional iron uptake systems, but expression among the core genome remained consistent (25).

We observed an increased motility phenotype in uropathogenic strain *E. coli* CFT073 under chemically induced iron-limited conditions. This phenotype corresponded to an increase in flagellin transcript, protein, and surface expression of flagella. Knowing that the well-studied Fur regulator was likely not facilitating this regulation (25), we took an unbiased forward genetic screening approach to identify the mediator of the observed iron-restricted enhanced motility phenotype. Through systematic deletion of regulatory genes, we ultimately found that deletion of the gene encoding dihydrolipoyl dehydrogenase, *lpdA*, led to a loss of iron-mediated motility in CFT073. This mutant also had a severe fitness defect in the murine urinary tract. Collectively, we demonstrated that deletion of a highly conserved metabolism-related gene alters motility, a urovirulence phenotype, and leads to a defect in the uropathogenesis property both *in vitro and in vivo*.

## Results

### The flagellar promoter is highly active under iron-restricted conditions

Previous RNAseq studies performed in clinical UPEC isolate HM7, revealed that flagella-related genes were significantly upregulated when the bacterium was cultured in M9 minimal medium when a gene essential in the production of its sole siderophore was mutated (26) (Table S1). We reasoned that this phenomenon may be occurring because the strain was somewhat iron depleted without its’ sole siderophore, forcing it to rely on less efficient iron transport systems. Strain HM7 is part of a more recent collection, isolated in 2009 from a young, healthy patient with cystitis (27, 28). We wanted to determine if this phenomenon was also relevant in clinical strain *E. coli* CFT073, isolated in 1990 from a patient with bacteriuria, acute pyelonephritis and bacteremia (17). However, unlike HM7, CFT073 has three siderophores (29) and would necessitate creating a triple mutant to re-create similar experimental conditions. Therefore, we decided to take a chemical approach and test conditions in an iron-depleted medium. First, we determined the concentration of the iron chelator 2,2 dipyridyl (dip) in LB medium that would restrict iron availability, but not substantially affect bacterial growth and survival (Fig. S1). We observed that 300 µM dip would serve as the ideal concentration for minimal growth inhibition and growth could be rescued with the addition of exogenous 300 µM FeCl_3_ (**Fig. 1A**). These conditions were utilized throughout this study.

**Figure 1.**
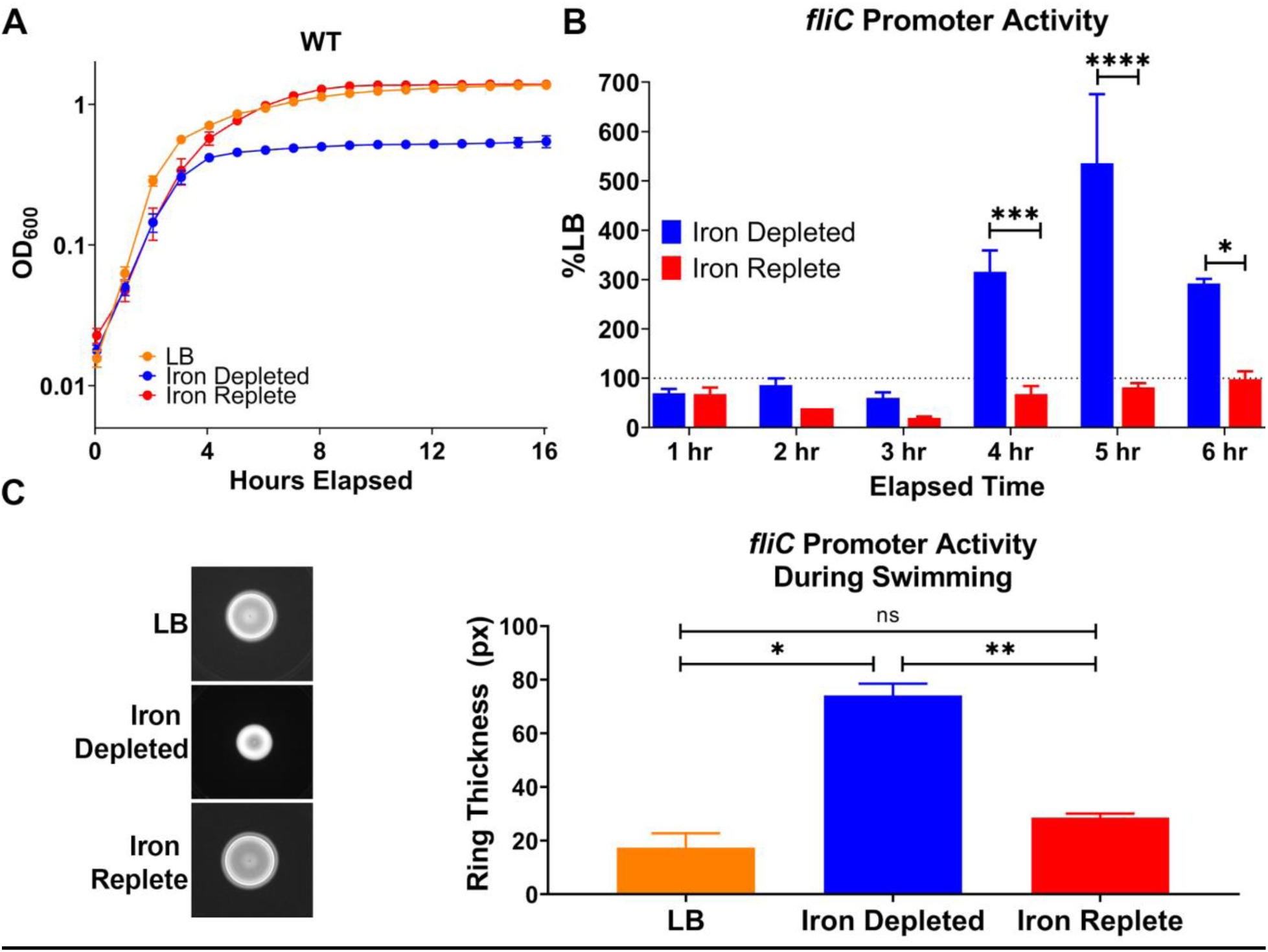
*fliC* promoter expression is elevated under iron-depleted conditions. (A) Uropathogenic *E. coli* CFT073 was cultured in LB, iron-depleted LB (supplemented with 300 μM 2,2 dipyridyl [dip], an iron chelator) and iron-replete LB (supplemented 300 μM dip and 300 μM FeCl_3_) 16 hours with aeration. Growth curves display averages from four biological replicates, error bars indicate ±SEM. (B) *fliC* promoter activity was assayed with a luciferase reporter over six hours in iron-deplete and iron-replete conditions and normalized to LB medium. Bars represent the mean from five biological replicates, error bars are ±SEM. Asterisks compare iron-depleted and iron-replete conditions, using 2way ANOVA with two-stage linear procedure of Benjamini Krieger and Yekutieli for multiple test corrections: * p< 0.05, *** p< 0.001, **** p<0.0001. (C) . *fliC* promoter activity was assessed during motility for 16 hours in semisoft agar at 37°C in LB, iron-depleted and iron-replete conditions. Representative image shows luciferase activity and the thickness of the outer ring was quantified in a bar graph. Bars display averages from four biological replicates, error bars indicate ±SEM. Asterisks compare iron-depleted and iron-replete LB medium, using 2way ANOVA Tukey’s multiple test corrections: * p< 0.05, ** p< 0.01.

Motility is an important process required for the ascension of UPEC into the upper urinary tract (1, 12). Lane *et al* (12) documented the kinetics and necessity of flagellar expression during active murine UTI utilizing a luciferase-expressing vector under the control of the native *fliC* promoter sequence. We used this vector to measure the activity and kinetics of the flagella subunit promoter under iron-depleted and iron-replete conditions, relative to unaltered LB medium. There were significant increases in *fliC* promoter activity at 4, 5, and 6 hpi (**Fig. 1B**). The largest difference between the iron-depleted and replete conditions was at 5 hpi, where there was over a five-fold increase in *fliC* promoter activation coupled with repression down to 81% of original levels in the iron-replete condition (**Fig. 1B**). Because promoter activity is the very first step in the central dogma of molecular biology, we wanted to assess if there was a commensurate change in the functional activity of the flagella. After overnight incubation, bacteria in iron-depleted LB swim agar moved a smaller distance in colony circumference compared to controls but demonstrated an increased intensity of *fliC* promoter activity (**Fig. 1C**). This *fliC* leading edge of the swim was quantified to be over four times thicker than the LB control, and this trend was repressed back to similar levels observed in the LB control by the addition of 300 µM FeCl_3_ exogenous iron in the presence of the iron chelator (**Fig. 1C**). These data demonstrated that CFT073, like observations made in another UPEC clinical isolate, upregulates flagella under iron-limited conditions.

### Flagella are upregulated at the gene, protein, and structural level

Gene expression studies performed in UPEC identified not only flagella machinery, but also chemotaxis components that were upregulated during iron deprivation (Table S1, (26)). Interestingly, the known flagellar master regulator complex, FlhDC (30, 31), was not found under these same conditions (Table S1). We therefore selected these specific genes to examine their expression during iron-depleted and -replete growth conditions, for validation. Similar to what was observed in other clinical UPEC strains, chemotaxis genes (*cheW* and *cheY*) and flagella machinery (*fliC* and *fliA*) were affected, but regulator *flhD* was not (**Fig. 2A**). The magnitude of upregulation was greatest in the flagellar genes and expression of all genes was able to be repressed when iron was added back to cultures (**Fig. 2A**). These trends were also observed at a protein level. Detectable FliC from bacteria cultured in iron-deprived conditions significantly increased over six-fold (**Fig. 2B**). Congruently, an increase in the average number of flagella per bacterial cell was observed and quantified via electron microscopy (**Fig. 2C**). A single flagellum would normally decorate the bacterium in LB or iron-replete conditions, while this number increased to two under iron depletion (**Fig. 2C**). These data, in conjunction with those in Figure 1, prove that the iron responsiveness of the flagella machinery in UPEC extends from promoter activity to the production of functional flagellar structures.

**Figure 2:**
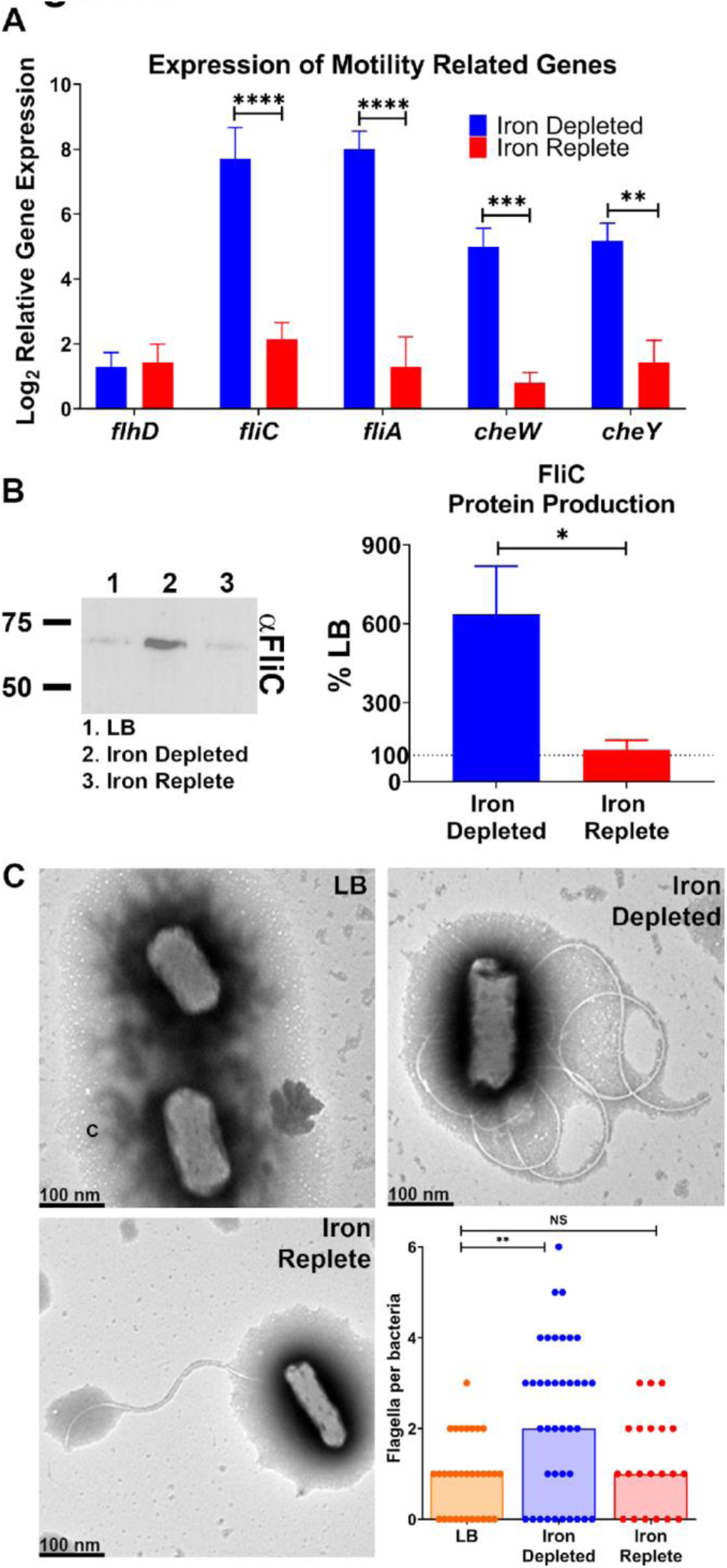
*fliC* gene expression and protein abundance are elevated under iron limitation, leading to an increase in surface-expressed flagella. CFT073 was cultured in LB, iron-depleted LB (supplemented 300 μM 2,2 dipyridyl [dip], an iron chelator) and iron-replete LB (300 μM dip+300 μM FeCl_3_) for five hours. (A) RNA was extracted and expression of indicated genes was assayed via qRT-PCR and compared to the LB control. n=4. (B) Cultures were normalized by OD_600_ and the whole cell lysate was immunoblotted. Densitometry of FliC was calculated, n=6, and representative immunoblot is shown. Bars indicate mean, error bars are ±SEM, *, p<0.05, ****p<0.0001, determined by paired *t*-test. (C) Representative electron micrographs of CFT073 cultured in either LB, iron-depleted or iron-replete conditions. Bacteria were fixed with 2.5% glutaraldehyde and stained with 1% phosphotungstic acid and imaged via TEM. The flagella per cell were quantified; each dot represents an individual bacterium. Bars represent the median, ** p< 0.01 determined by Kruskal-Wallis test with Dunnet’s multiple comparison.

### Global regulator mutants *fis*, *crp*, and *arcA* respond differently to iron replete conditions

Both motility-based and iron responsiveness regulons have been thoroughly studied in *E. coli*. FlhDC is well-known as the master regulator of the flagellar operon (32), but this system was not upregulated in response to iron depletion (25). To take an unbiased and global approach in defining the CFT073 iron-mediated flagella upregulation, we generated deletions in eight master transcriptional regulators (33). None of these mutants exhibited drastically different growth profiles under iron-depleted and -replete conditions (**Fig. S2**). We then examined the swimming ability of these mutants to select candidates for follow-up (**Fig. 3A**). Interestingly, almost every single mutant had a significant change in swimming motility. The Δ*fis,* Δ*crp,* Δ*arcA* and Δ*hns* mutants either had reduced motility or were non-motile, while the Δ*lrp,* Δ*ihf*, and Δ*fnr* mutants were hyper-motile when compared to WT (**Fig. 3A**). Δ*narL* was the only mutant with no change in motility. We were encouraged by these results and went on to specifically assess the loss of iron-regulated flagellin expression in these mutants (**Fig. 3B, C**). To do so, we compared the expression of *fliC* transcript in both the iron-depleted and -replete conditions of the mutants to WT. If the mutant had over a 2-fold change in expression when compared to WT in either condition, we marked it for further study (**Fig. 3B, C**). Based on these criteria, Δ*fis,* Δ*crp*, Δ*arcA*, and Δ*hns* lost the ability to regulate *fliC* in an iron-dependent manner (**Fig. 3B**, **C**). However, Δ*hns* had nearly undetectable levels *of fliC* via qRT-PCR (CT value >30) in any condition and was therefore eliminated from follow up analyses. We examined the direct regulons of Δ*fis,* Δ*crp*, and Δ*arcA* to narrow down the potential iron-mediated motility mechanism in UPEC. Each of these master regulators is known to influence several hundred genes (33); however, only five genes were present in all three regulons (**Fig. 3D**). These were *acnB*, *ptsG*, *xylA*, *lpdA*, and *fumB*. Four of these genes are involved in metabolism-related processes and three of them have metallic cation binding sites (*fumB*, *acnB*, *xylA*).

**Figure 3.**
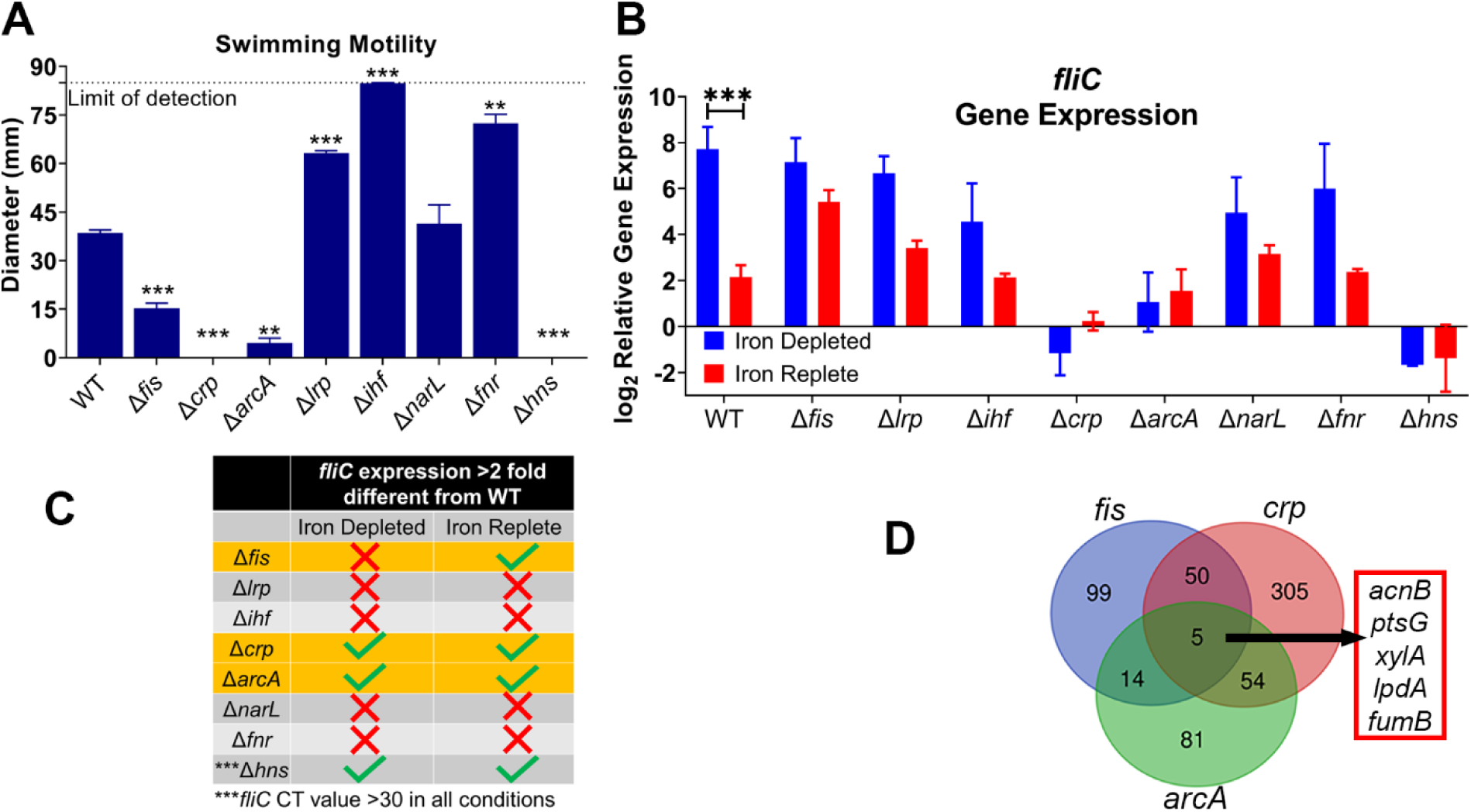
Loss of *fis, crp,* or *arcA* disrupts iron-mediated *fliC* expression. Master regulator mutants were generated using λ red mutagenesis. Strains were cultured for five hours in LB, iron-depleted (LB supplemented with 300 μM dip) and iron-replete (LB supplemented with 300 μM dip and 300 μM FeCl_3_). (A) *fliC* expression was assessed and normalized to LB as in previous assays. n=4, bars indicate mean, error bars are ±SEM, ***p<0.001, determined by 2way ANOVA with Sidak’s multiple comparison test. (B) Motility was assessed in semisoft agar at 30°C, n=4. ** p<0.01, ***p<0.001, RM ANOVA, Dunnet’s correction. (C) Strategy to cull transcription factor hits. (D) Genes regulated by *fis, crp* and *arcA*.

### Mutation of *lpdA* encoding lipoamide dehydrogenase causes loss of iron-regulated *fliC* **expression and motility.**

To further pursue the regulatory mechanism of iron-mediated motility, we generated knockouts in each of the previously determined genes of interest and examined the expression of *fliC* under iron limitation and the swimming motility of these mutants (**Fig. 4A**). The Δ*lpdA* mutant was the only construct that was non-motile in swim agar (**Fig. 4A**). Similarly, we found that the Δ*lpdA* mutant was the only strain where *fliC* expression was no longer regulated by iron (**Fig. 4B**). Even with these two defects, the mutant was able to grow similarly to wild-type under iron-restricted conditions in LB medium (**Fig. S3**). LpdA is the E3 component of the pyruvate dehydrogenase complex, which converts pyruvate into acetyl-CoA that feeds into the TCA cycle, connecting glycolysis with the TCA cycle under aerobic conditions (34, 35). Specifically, LpdA binds the coenzyme FAD which facilitates the oxidation of dihydrolipoate back to lipopate, producing FADH_2_ in the process (34, 35). Loss of LpdA results in accumulation of intracellular pyruvate and glucose (34). However, *E. coli* is still able to use glucose as a sole carbon source in the absence of LpdA due to another enzyme, PoxB, that can directly convert pyruvate into CO_2_ and acetate (34).

**Figure 4.**
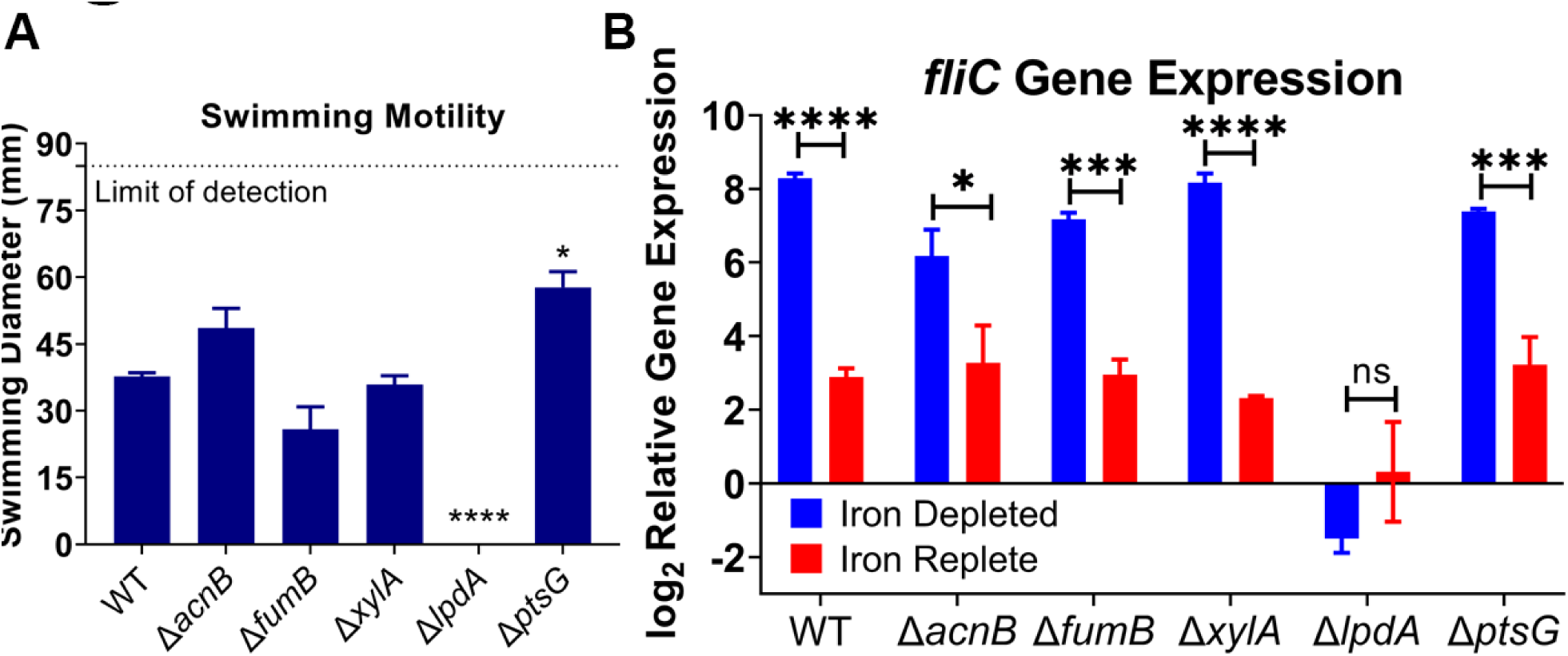
Loss of *lpdA* disrupts iron-mediated *fliC* response. Mutants were generated using λ red mutagenesis. Strains were cultured for five hours in LB, iron-depleted (LB supplemented with 300 μM dip) and iron-replete (LB supplemented with 300 μM dip and 300 μM FeCl_3_). (A) Motility was assessed in semisoft agar at 30°C, n=6 *p<0.05, ****p<0.0001, RM ANOVA, Dunnet’s correction. (B) *fliC* expression levels were assessed five hours post inoculation and normalized to LB as in previous assays. n=4, Bars represent the mean, error bars are ±SEM, * p<0.05, ***p<0.001, ****p<0.0001 determined by 2way ANOVA with Sidak’s multiple comparison test.

After selecting Δ*lpdA* as our gene of interest, we continued to explore the mutant strain within the context of iron-depletion. Knowing that gene expression of *fliC* was unresponsive (**Fig. 4B**), we turned our attention to potential deficiencies at the protein and structural assembly levels, and finally, uropathogenesis. Δ*lpdA* produced about two-fold more FliC protein in iron-depleted LB compared to LB (**Fig. 5A**), but the differences observed in iron-depleted and -replete conditions were not significant. This suggests a decreased sensitivity of an iron-based response. The Δ*lpdA* mutant produced almost no flagella per bacterial cell; the median number of flagella in all three tested conditions was zero (**Fig. 5B, Fig. S4A**). However, the iron-depleted condition had a higher range of flagella, with a maximum of five flagella per bacterial cell, compared to zero under iron-replete conditions (**Fig. S4B**). Furthermore, these values are significantly different from the WT values, (**Fig. 5B**) reflecting the changes seen via western blot and indicating a general dysregulation of flagellar production.

**Figure 5.**
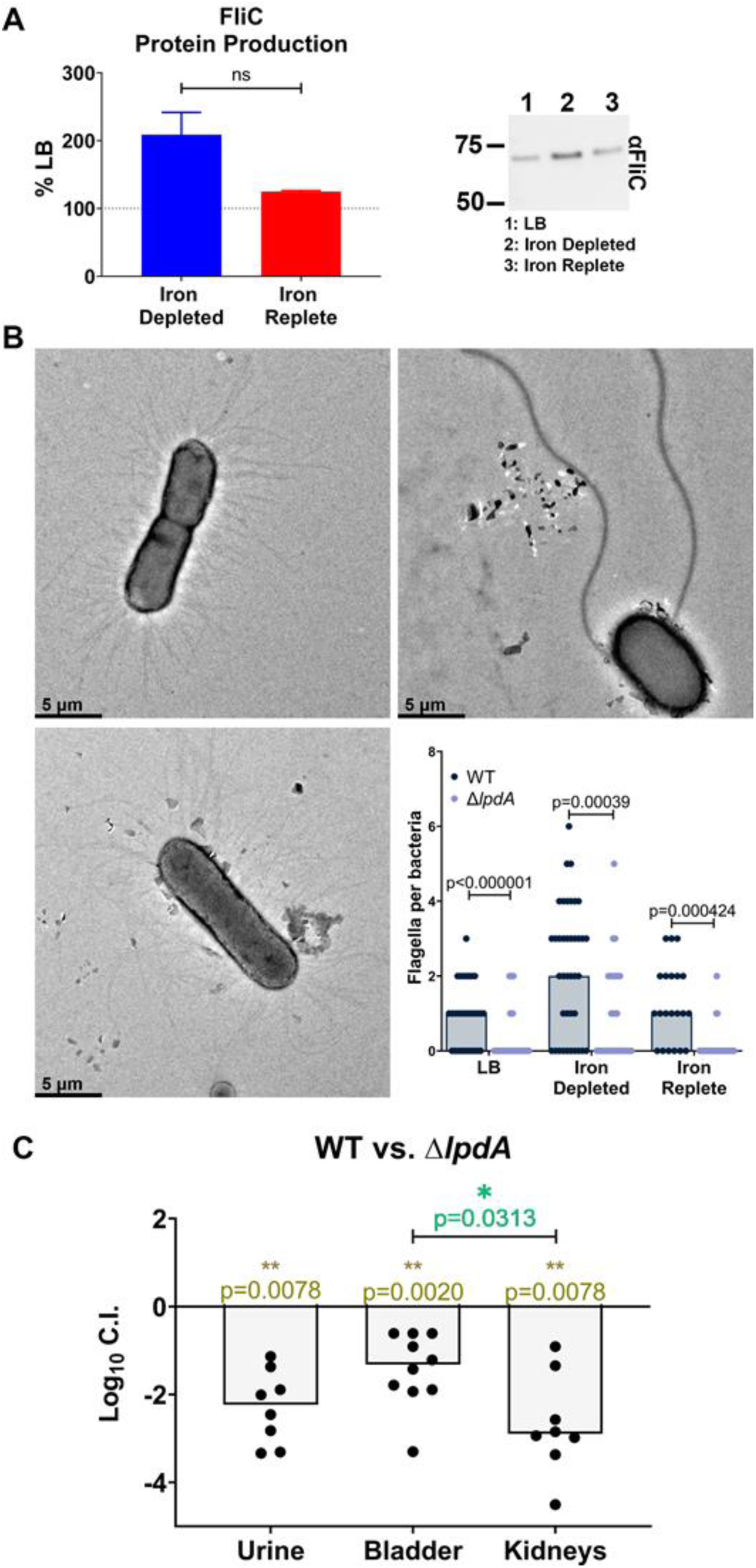
Loss of lpdA attenuates FliC protein production and flagella production under iron limitation, resulting in host fitness defects. (A) Cultures were normalized by OD_600_ and the whole cell lysate was immunoblotted. Densitometry of FliC was calculated, n=4, and representative immunoblot is shown. Bars represent the mean, error bars are ±SEM, not significant (ns) determined by unpaired *t*-test. (B) Representative electron micrographs of CFT073 cultured in either LB, iron-depleted or iron-replete conditions. Bacteria were fixed with 2.5% glutaraldehyde and stained with 1% phosphotungstic acid and imaged via TEM. The number of flagella per bacterium was quantified; each dot represents an individual bacterium. Asterisks compare WT and mutant constructs in LB, iron-depleted or iron-replete conditions using Mann-Whitney tests with a multiple test correction using FDR. (C) WT CFT073 and the *lpdA* mutant were combined in a 1:1 ratio and transurethrally inoculated into the bladders of CBA/J mice. Bars represent the median, while each dot represents an individual animal, n=10. Significance was determined using Wilcoxon’s signed-rank test with indicated *p* values, and burden between bladder and kidneys compared using Wilcoxon’s test.

### Surface-expressed flagella are reduced in the *lpdA* mutant but their expression can be genetically complemented

We have confirmed that the loss of *lpdA* diminishes the iron-mediated response of flagellar upregulation (**Fig. 4B, 5A, B**). In fact, the Δ*lpdA* mutant has significantly reduced flagella expression leading to a loss of motility under all conditions tested (**Fig. 4A**). To confirm that these phenomena were a direct result of *lpdA*, we complemented *in trans* under the control of its native promoter and tested resulting phenotypes. The motility of the mutant was restored to the level of WT with the addition of the *lpdA* vector compared to the empty vector negative control (**Fig. S5A**). We were also able to rescue flagella production in both the LB and iron-deplete conditions in the genetically complemented strain (**Fig. S5B, C**). We did not observe rescue in the iron-replete condition (**Fig. S5D**); however, flagella production of the WT strain harboring the empty vector was also dysregulated when compared to WT alone, which may account for this result. Ultimately, using a systematic approach, we determined that loss of *lpdA* causes a lack of iron concentration-dependent motility responsiveness, and an overall loss of flagella expression.

Finally, we wanted to assess the effect that *lpdA* might have on pathogenesis. We transurethrally inoculated female CBA mice with equal ratios of WT and Δ*lpdA* bacteria (total of 2×10^8^ cfu) using the murine model of ascending UTI (36). Loss of *lpdA* in CFT073 resulted in over a 20-fold fitness defect in the bladder and a 500-fold defect in the kidneys when co-challenged with the wild-type parent strain (**Fig. 5C**); The greater defect in the kidneys is unsurprising given Δ*lpdA* is impaired in swimming motility. This further underscores the importance of swimming motility for kidney infection as well as the appropriate response to iron levels. Altogether, these results show the importance of *lpdA* and the iron-regulated motility for the full pathogenesis of UPEC.

## Discussion

Most iron within the human host is sequestered by specific host proteins such as transferrin to control bacterial growth (24). In response, pathogenic bacteria employ numerous well-studied strategies to acquire iron from these sequestered sources including siderophore production and other iron import systems (37). UPEC, the main causative agent of uncomplicated UTI, is no exception and uses these various systems to survive within the urinary tract. Indeed, iron acquisition is so crucial to infection and pathogenesis that antigens from these systems have been used in trial vaccines against UPEC (38, 39) due to their increased surface expression during infection.

Motility is another highly studied arm of UPEC virulence; it enables bacteria to ascend the ureters to the kidneys more effectively and rapidly (12). Despite the significance of these two virulence traits, to date, there are no studies linking these two virulence mechanisms in UPEC. Here, we demonstrate that an iron-restricted environment leads to an increase in motility-related functions, spanning the regulatory dogma, from gene expression of flagellar components to an increase of flagella on the surface of the bacteria.

We observed genetic regulation between iron levels and motility in a clinical isolate from 2009 (27, 28). To ensure this was a universal response in UPEC, we tested this response in uropathogenic type strain *E. coli* CFT073 under iron-limited conditions using chemical iron chelation. *fliC* promoter activity, transcript and protein production were all significantly increased under iron limitation (**Fig. 1B**, **Fig. 2A, B**). This response extended to functional consequences; strain CFT073 cultured under iron-depleted conditions was decorated with more flagella (**Fig. 2C**) and the leading edge of swimming bacteria expressed *fliC* more strongly (**Fig. 1C**). The addition of exogenous iron was able to re-repress all responses (**Fig 1, 2**) demonstrating that these are iron-specific responses and not attributable to off-target effects of the iron chelator. Altogether, these results show a robust and consistent increase in motility and motility-related systems when UPEC is subjected to iron limitation. Perhaps an iron-limited environment could serve as one of many cues to the bacteria upon entering the host and thus need to upregulate flagellar machinery. We consider this despite there being no known chemoreceptor for iron in these bacteria.

It is interesting to speculate that iron-mediated motility in *E. coli* has remained understudied because the seminal papers that established conditions promoting motility used commensal isolate K12 and observations and conclusions drawn from a non-pathogenic isolate are not always applicable to a pathogenic strain. For example, while the motility of K12 is inhibited at 37°C (15, 40) our study shows robust *fliC* expression and upregulation in UPEC strain CFT073 at this temperature (**Fig. 1C, 2A**) as well as an ability to swim (**Fig. 1C**). Furthermore, while our study reveals a strong, reproducible response where iron limitation induces *fliC* expression in CFT073, the opposite has been shown in K12, actually increasing iron concentration elevated *fliC* expression in a K12 derivative (41). This starkly emphasizes the differential regulatory responses between commensal and pathogenic isolates. Perhaps the iron-regulated motility response could be used as a distinguishing feature of uropathogens. Or the response could be characteristic of pathogenic *E. coli* that infect other body sites, such as the bloodstream or gut. Determining the conservation of this response mechanism among UPEC and other uropathogens is a future research direction.

After establishing the consistency of the iron-regulated motility response, our next objective was to define the underlying mechanism. Two widely studied iron regulators are Fur (Ferric Uptake Regulator) (42, 43) and RyhB (a small RNA) (44). However, a recent paper (25) defined both the direct and indirect regulons of Fur and RyhB in CFT073 and not a single component of the flagellar machinery was uncovered. Therefore, we decided to investigate eight master regulators (*fis, lrp, ihf, crp, arcA, narL, fnr,* and *hns*) and determine if they were involved in iron-mediated motility (**Fig. 3**). We continued this investigation to determine the downstream cascade of signaling leading to *fliC* upregulation. Loss of *fis, crp* or *arcA* resulted in dysregulation of *fliC* gene expression in iron-depleted conditions (**Fig. 3B**). These three regulators shared five genes in their direct regulons: *acnB, ptsG, xylA, lpdA,* and *fumB*. We reasoned that these genes might have a more direct role in regulating iron-mediated motility, and indeed the *lpdA* mutant resulted in dysregulation of iron-mediated motility (**Fig 4B, 5A, B**). While this mutant was non-motile, it still produced flagella that reached the bacterial surface, albeit to a lesser degree (**Fig. 5A, B**). Lastly, we assessed the fitness of the *lpdA* mutant in the murine model of UTI. The mutant had a significant and severe defect in all three organ sites tested (urine, bladder and kidneys), with the most pronounced defect in the kidneys (>100 fold disadvantage, **Fig. 5C**). We were surprised that *lpdA* appears to be integral to iron-mediated motility, given that it has no known iron-binding site. Previous studies have investigated the role that *lpdA* plays in metabolism; it encodes a part of the pyruvate dehydrogenase complex that connects glycolysis with the TCA cycle (35). However, it is worth noting that there are functionally redundant enzymes to LpdA such as PoxB (45) that can compensate for the absence of LpdA, enabling a *lpdA* mutant to utilize glucose as a sole carbon source. Therefore, we believe the loss of fitness during murine infection is not solely due to metabolic dysfunction. Moreover, previous papers (46, 47) have shown that glycolysis is dispensable for UPEC during UTI. This indicates that our observed defect could be directly attributable to the loss of motility, ultimately leading to decreased murine colonization.

It was interesting to find that a metabolic gene would so drastically affect flagella-mediated motility, and it is exciting to speculate on the mechanism driving this result. Loss of LpdA results in increased intracellular glucose (34), and it has been established that glucose suppresses production of flagella in non-pathogenic *E. coli* (40). While the uropathogenic *lpdA* mutant still expresses a few surface flagella, it is possible that the buildup of intracellular glucose levels results in the reduction of motility. Perhaps, future studies should include the addition of glucose to the same growth conditions to chemically suppress the iron-regulated expression of *fliC,* thus phenocopy the *lpdA* mutant.

Altogether, our study highlights the possible distinctions between uropathogenic bacteria and their commensal counterparts. This work further emphasizes the importance of studying differential regulatory networks in pathogens versus commensals, even among the same bacterial species. We discovered a novel regulatory pathway in UPEC that leads to the upregulation of *fliC* and surface-expressed flagella, that is specifically deployed in an iron-depleted environment. This contributes to our biological understanding of the myriad of host virulence mechanisms and how they contribute to disease. Linking motility and iron responsiveness, two highly studied virulence mechanisms, opens new avenues of research for this ubiquitous pathogen and the findings there can potentially be applied to other pathogenic strains of *E. coli*, or other uropathogens.

## Materials and Methods

### Bacterial growth and culture

*E. coli* CFT073 was routinely cultured overnight in Luria broth (LB) with aeration at 37°C, unless otherwise stated. Cultures were inoculated with a single isolated colony. Kanamycin (25 µg/mL) was added to the medium for designated mutant constructs, and ampicillin (100 ug/mL) was added for maintenance of plasmids. Strains used are in **Table 1**.

**Table 1:**
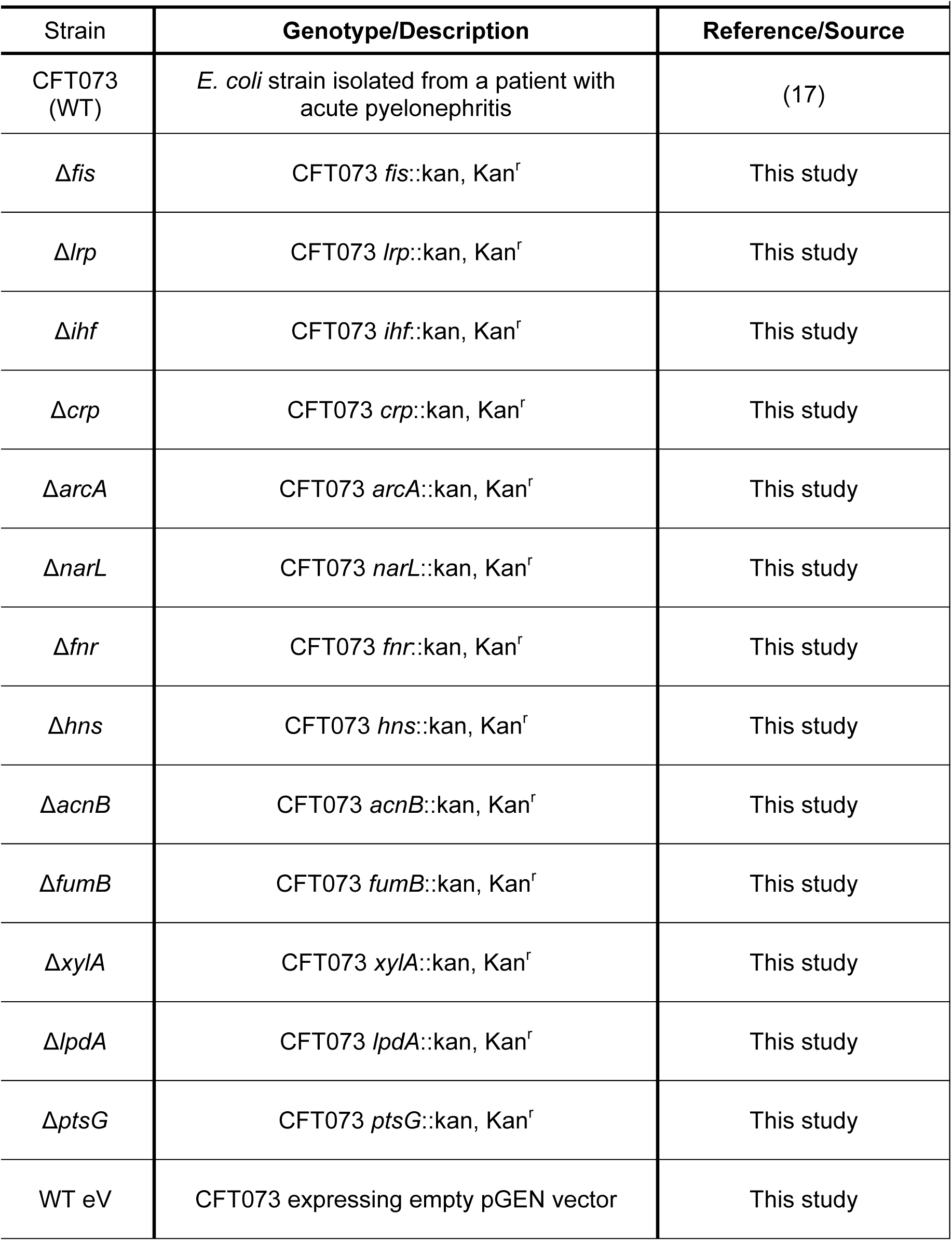
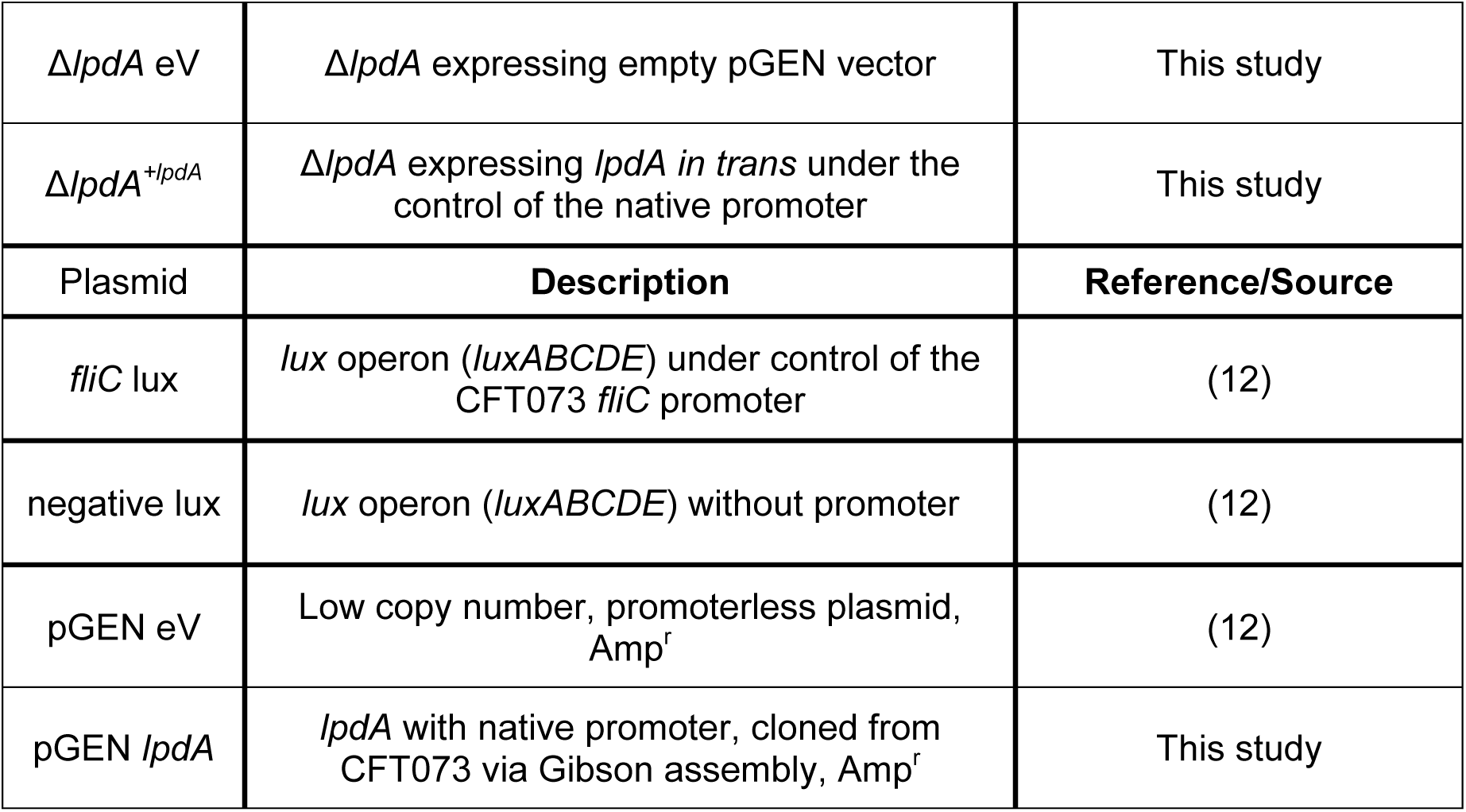
List of Strains and Plasmids.

### Growth curves

Strains were cultured overnight in LB with appropriate antibiotics at 37°C with aeration. The next day, cultures were back-diluted 1:100 into indicated conditions and incubated with shaking at 37°C for 16 hours using the Bioscreen-C automatic growth curve analyzer.

### Bacterial mutants and genetic complementation

The lambda red recombinase system was used to generate mutants in CFT073 (48). Homology regions of 35 bp, both upstream and downstream of the gene of interest, were used to flank the kanamycin resistance cassette amplified from pKD4. This PCR product was introduced into wild-type strain carrying the recombinase expression system encoded on pKD46 (48). Complementation vectors in pGEN were constructed using Gibson Assembly. Primers used are in **Table 2**.

**Table 2:**
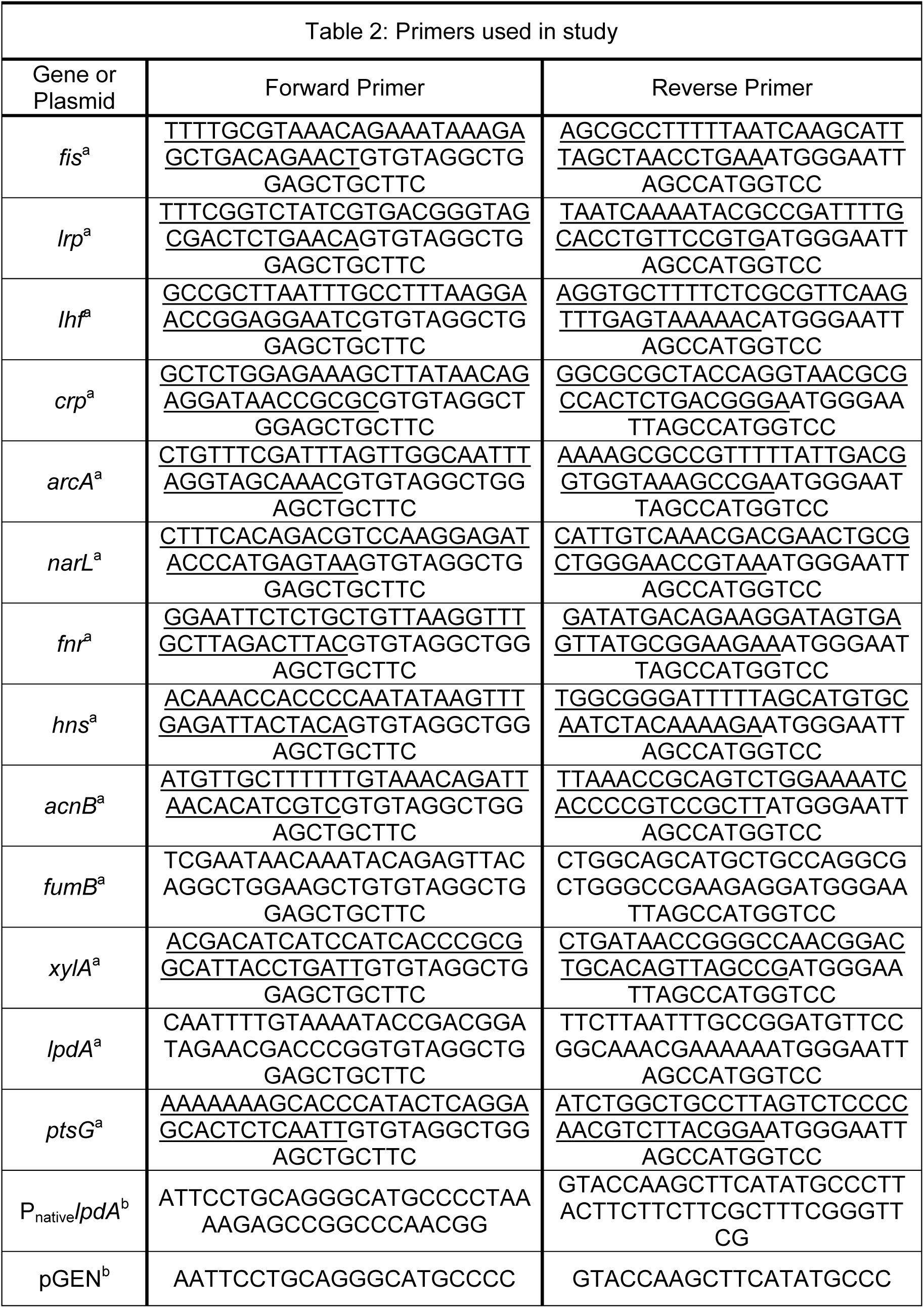
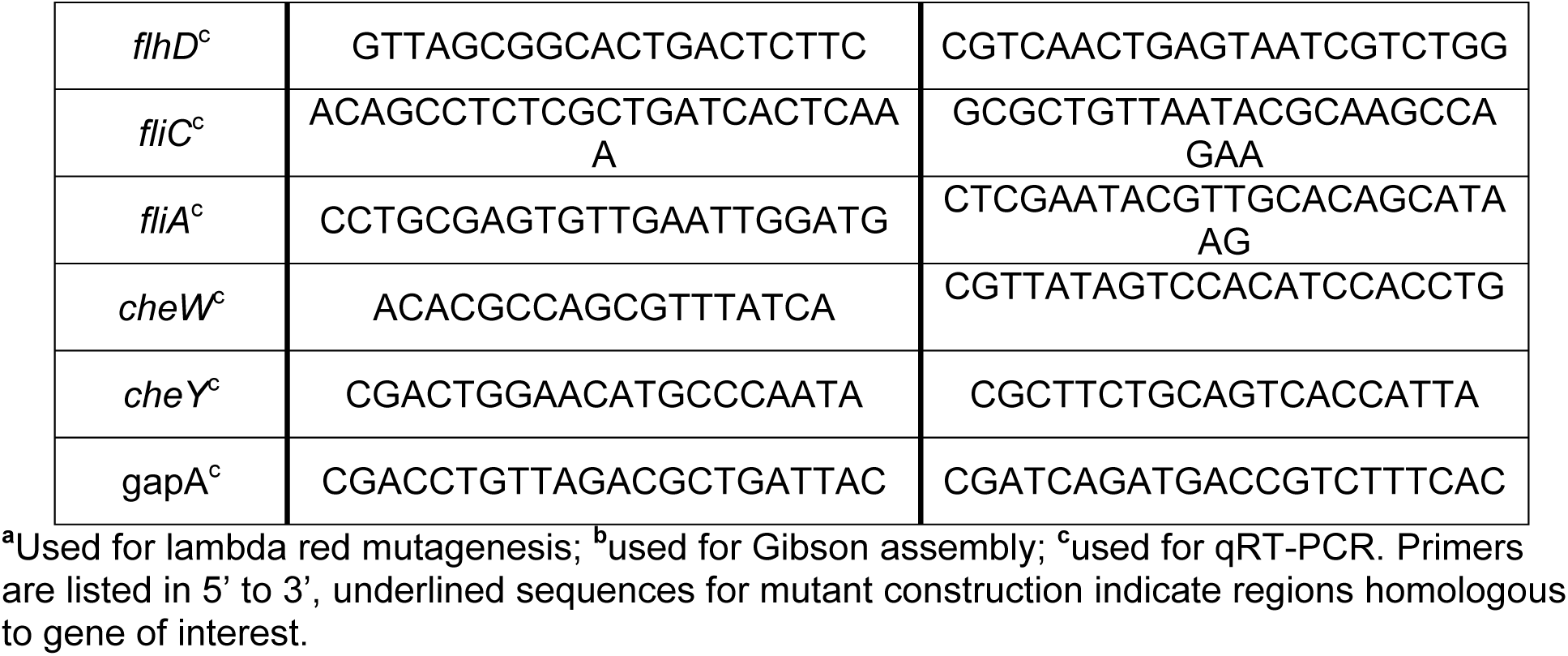
Primers used in study.

### *fliC* promoter luminescence activity

We used two reporter constructs that have been previously published (12). One, where the CFT073 *fliC* promoter-controlled luciferase production (*luxABCDE*), and a promoter-less negative control. We cultured both strains overnight with appropriate antibiotics at 37°C with aeration. The next morning, cultures were back-diluted 1:100 into LB, LB supplemented with 300 μM dip (iron-deplete), and LB supplemented with 300 μM dip and 300 μM FeCl_3_ (iron-replete) and cultured at 37°C with aeration. Cultures were sampled every hour for six hours for growth (via OD_600_) and luminescence. Luminescence was quantified with a Synergy H1 reader (Agilent BioTek) using a black-sided 96-well plate. Luminescence was normalized by colony forming units (CFU) at an OD_600_ reading of 1=7.5×10^8^ CFU/mL, the background luminescence calculated by the values of the negative control and subtracted from the experimental group. Finally, these normalized luminescence values were reported per 10^6^ CFU.

### Swimming assay

CFT073 motility was assessed in semi-soft swimming agar, formulated as previously described (49). Briefly, after overnight growth, bacterial cultures were normalized to an OD_600_=1 in 10 mM HEPES (pH 8.4). Using a sterile inoculating needle, bacteria were introduced to the center of the agar. Plates were incubated for 16 hr at 30°C. Swim diameters were measured and recorded in mm.

### Luminescent swimming assay

Strains harboring the *fliC* reporter plasmid were cultured overnight under antibiotic selection. The next morning, cultures were normalized as described in the swimming assay. These cultures were inoculated into semi-soft (0.25% agar) LB agar and incubated for 16 hours at 37°C. The next morning, plates were imaged on a Biorad imager using the chemiluminescent channel to visualize activity of the *fliC* promoter. The bright outer ring, representing the actively swimming bacteria, was measured at its thickest point.

### Quantitative Reverse-Transcriptase PCR

qRT-PCR was conducted as previously published (26). Briefly, strains were cultured overnight in LB with appropriate antibiotics at 37°C with aeration. The next morning cultures were back-diluted 1:100 into LB medium as well as the iron-deplete and iron-replete conditions. Strains were cultured for five hours at 37°C with aeration before harvest and treatment with bacterial RNAprotect (Qiagen). Bacterial pellets were stored at -80°C until RNA isolation.

RNA was isolated using Qiagen’s RNAeasy kit. Genomic DNA was removed with Turbo DNAse free kit, and then RNA was converted into cDNA using Biorad’s iscript. qRT-PCR reactions were performed on 10 ng of cDNA in technical duplicate using SyberGreen reagent and the Quantstudio3 (Applied Biosystems). Gene expression was calculated using the 2^-ΔΔCT^ method with *gapA* as a housekeeping gene (26) and compared to the LB only condition. Primers used are in **Table 2**.

### Western Blot

Strains were cultured as described for qRT-PCR. Cultures were then normalized to an OD_600_ of 0.4, pelleted at low speed (<5,000 RPM), resuspended in SDS-loading buffer and boiled for five minutes. Proteins were separated on a 4-20% SDS-PAGE gel (Biorad) and transferred to nitrocellulose. Membranes were probed with αH1 FliC antibody (Rockland), using a α rabbit secondary antibody conjugated to HRP.

### Electron microscopy

Strains were cultured as previously described. A drop of bacterial culture was applied to glow-discharged carbon coated copper grids with a 400 mesh (EMS) and allowed to incubate for three minutes. Excess culture was wicked away and a drop of 2.5% glutaraldehyde (EMS) was applied to the grids for five minutes to fix bacteria. Grids were then washed by being plunged in water ten times and the excess water wicked away. A 1% phosotungistic acid stain was applied to the grids for 30 seconds to stain the samples and the excess aspirated away to dry the grids. Samples were imaged using a Morgangi (FEI, Hillsboro, OR) operated at an acceleration voltage of 100 kV and equipped with a 1kx1k charge-coupled-device (CCD) camera (ATM) at 1,800x magnification.

### Murine model of ascending UTI

Mice were inoculated as previously published (36). Briefly, female CBA/J mice were transurethrally inoculated with 10^8^ CFU of 1:1 mixed wild-type and *lpdA* mutant of CFT073. After 48 h, urine, bladder, and kidneys were harvested to enumerate bacterial burden. Differential plating on Luria agar plates containing antibiotics to quantify the wild-type to mutant ratio. The ratio of strains coming out of the infection was compared to the ratio of the inoculum going into the infection to determine a competitive index (26).

## Acknowledgments

We would like to acknowledge Dr. Melanie Pearson for her thoughtful discussion. We would also like to acknowledge our sources of funding from the National Institutes of Health: R01AI059722, R01AI165582, and F32AI147527 as well as the National Science Foundation: DGE1841052. We acknowledge the use of the U-M LSI cryo-EM facility, and University of Michigan LSI IT support. We thank University of Michigan BSI and LSI for significant support of the cryo-EM facility.

## Supplemental Figure Legends

**Figure S1. Growth of *E. coli* CFT073 in LB medium supplemented with increasing concentrations of iron chelator 2,2’-dipyridyl (dip).** CFT073 was cultured overnight in LB, and then subcultured 1:100 and incubated at 37°C with aeration in LB supplemented with indicated concentrations of 2,2 dipyridyl (dip), an iron chelator. OD_600_ was measured on a plate reader for 16 hours, taking a reading every hour. Results are an average of two biological replicates; error bars represent ±SEM.

**Figure S2. Growth of WT *E. coli* CFT073 and indicated mutants in LB, iron-depleted and iron-replete conditions.** The iron-depleted condition is LB supplemented 300 μM 2,2 dipyridyl (dip), an iron chelator while iron-replete is LB supplemented 300 μM 2,2 dipyridyl (dip) and 300 μM FeCl_3_. Strains were cultured overnight in LB with antibiotics where appropriate and then subcultured 1:100 and incubated in the indicated conditions at 37°C with aeration. OD_600_ was measured on a plate reader for 16 hours, taking a reading every hour. Results are the average of four to five biological replicates; error bars represent ±SEM. WT growth curves are the same as the one shown in Fig. 1A.

**Figure S3. Growth of WT *E. coli* CFT073 and indicated mutants in LB, iron-depleted and iron-replete conditions.** The Iron-depleted condition is LB supplemented 300 μM 2,2 dipyridyl (dip), an iron chelator while iron-replete is LB supplemented 300 μM 2,2 dipyridyl (dip) and 300 μM FeCl_3_. Strains were cultured overnight in LB with antibiotics where appropriate and then subcultured 1:100 and incubated in the indicated conditions at 37°C with aeration. OD_600_ was measured on a plate reader for 16 hours, taking a reading every hour. Results are an average of four biological replicates; error bars represent ±SEM.

**Figure S4. Flagella per bacterial cell in a *lpdA* mutant.** (A) Flagella per bacterium were quantified in each of the indicated conditions; each dot represents an individual bacterium. (B) The quartiles of flagellar number under each indicated condition.

**Figure S5. Complementation of the *lpdA* mutant.** The *lpdA* mutant was complemented *in trans* using the pGEN vector under control of the native *lpdA* promoter. WT eV is the WT strain expressing an empty vector; Δ*lpdA* eV is the mutant expressing an empty vector while Δ*lpdA^+lpdA^*is the complemented strain. These three constructs were tested for their (A) motility, and flagellar production in (B) LB, (C) iron depleted, and (D) iron replete conditions.

## References

1. Bacheller CD, Bernstein JM. 1997. URINARY TRACT INFECTIONS. Medical Clinics of North America 81:719–730.

2. Geerlings Suzanne E. 2016. Clinical Presentations and Epidemiology of Urinary Tract Infections. Microbiology Spectrum 4:10.1128/microbiolspec.uti-0002-2012.

3. Foxman B. 2010. The epidemiology of urinary tract infection. Nature reviews Urology 7:653–660.

4. Flores-Mireles AL, Walker JN, Caparon M, Hultgren SJ. 2015. Urinary tract infections: epidemiology, mechanisms of infection and treatment options. Nature Reviews Microbiology 13:269–284.

5. Foxman B. 2014. Urinary tract infection syndromes: occurrence, recurrence, bacteriology, risk factors, and disease burden. Infect Dis Clin North Am 28:1–13.

6. Stamm WE, Hooton TM. 1993. Management of Urinary Tract Infections in Adults. New England Journal of Medicine 329:1328–1334.

7. Schwartz L, de Dios Ruiz-Rosado J, Stonebrook E, Becknell B, Spencer JD. 2023. Uropathogen and host responses in pyelonephritis. Nature Reviews Nephrology doi:10.1038/s41581-023-00737-6.

8. Lane MC, Mobley HLT. 2007. Role of P-fimbrial-mediated adherence in pyelonephritis and persistence of uropathogenic Escherichia coli (UPEC) in the mammalian kidney. Kidney International 72:19–25.

9. Smith Sara N, Hagan Erin C, Lane MC, Mobley Harry LT. 2010. Dissemination and Systemic Colonization of Uropathogenic Escherichia coli in a Murine Model of Bacteremia. mBio 1:10.1128/mbio.00262-10.

10. Wagenlehner FME, Lichtenstern C, Rolfes C, Mayer K, Uhle F, Weidner W, Weigand MA. 2013. Diagnosis and management for urosepsis. International Journal of Urology 20:963–970.

11. Little P, Merriman R, Turner S, Rumsby K, Warner G, Lowes JA, Smith H, Hawke C, Leydon G, Mullee M, Moore MV. 2010. Presentation, pattern, and natural course of severe symptoms, and role of antibiotics and antibiotic resistance among patients presenting with suspected uncomplicated urinary tract infection in primary care: observational study. BMJ 340:b5633.

12. Lane MC, Alteri CJ, Smith SN, Mobley HLT. 2007. Expression of flagella is coincident with uropathogenic *Escherichia coli* ascension to the upper urinary tract. Proceedings of the National Academy of Sciences 104:16669–16674.

13. Thormann KM, Beta C, Kühn MJ. 2022. Wrapped Up: The Motility of Polarly Flagellated Bacteria. Annu Rev Microbiol 76:349–367.

14. Adler J. 1966. Chemotaxis in bacteria. Science 153:708–16.

15. Morrison RB, McCapra J. 1961. Flagellar Changes in *Escherichia coli* induced by Temperature of the Environment. Nature 192:774–776.

16. Yang W, Briegel A. 2020. Diversity of Bacterial Chemosensory Arrays. Trends in Microbiology 28:68–80.

17. Welch RA, Burland V, Plunkett G, Redford P, Roesch P, Rasko D, Buckles EL, Liou S-R, Boutin A, Hackett J, Stroud D, Mayhew GF, Rose DJ, Zhou S, Schwartz DC, Perna NT, Mobley HLT, Donnenberg MS, Blattner FR. 2002. Extensive mosaic structure revealed by the complete genome sequence of uropathogenic *Escherichia coli*. Proceedings of the National Academy of Sciences 99:17020–17024.

18. Kaper JB, Nataro JP, Mobley HLT. 2004. Pathogenic *Escherichia coli*. Nature Reviews Microbiology 2:123–140.

19. Kondoh H, Ball CB, Adler J. 1979. Identification of a methyl-accepting chemotaxis protein for the ribose and galactose chemoreceptors of *Escherichia coli*. Proc Natl Acad Sci U S A 76:260–4.

20. Manson MD, Blank V, Brade G, Higgins CF. 1986. Peptide chemotaxis in *E. coli* involves the Tap signal transducer and the dipeptide permease. Nature 321:253–6.

21. Lane MC, Lloyd AL, Markyvech TA, Hagan EC, Mobley HL. 2006. Uropathogenic *Escherichia coli* strains generally lack functional Trg and Tap chemoreceptors found in the majority of *E. coli* strains strictly residing in the gut. J Bacteriol 188:5618–25.

22. Spurbeck Rachel R, Dinh Paul C, Walk Seth T, Stapleton Ann E, Hooton Thomas M, Nolan Lisa K, Kim Kwang S, Johnson James R, Mobley Harry LT, Bäumler AJ. 2012. *Escherichia coli* Isolates That Carry *vat*, *fyuA*, *chuA*, and *yfcV* Efficiently Colonize the Urinary Tract. Infection and Immunity 80:4115–4122.

23. Brumbaugh AR, Smith SN, Mobley HLT. 2013. Immunization with the Yersiniabactin Receptor, FyuA, Protects against Pyelonephritis in a Murine Model of Urinary Tract Infection. Infection and Immunity 81:3309–3316.

24. Murdoch CC, Skaar EP. 2022. Nutritional immunity: the battle for nutrient metals at the host–pathogen interface. Nature Reviews Microbiology 20:657–670.

25. Banerjee R, Weisenhorn E, Schwartz Kevin J, Myers Kevin S, Glasner Jeremy D, Perna Nicole T, Coon Joshua J, Welch Rodney A, Kiley Patricia J, Gottesman S. Tailoring a Global Iron Regulon to a Uropathogen. mBio 11:e00351–20.

26. Frick-Cheng AE, Sintsova A, Smith SN, Pirani A, Snitkin ES, Mobley HLT, Comstock LE. 2022. Ferric Citrate Uptake Is a Virulence Factor in Uropathogenic Escherichia coli. mBio 13:e01035–22.

27. Sintsova A, Frick-Cheng A, Smith S, Pirani A, Subashchandrabose S, Snitkin E, Mobley HLT. 2019. Genetically diverse uropathogenic *Escherichia coli* adopt a common transcriptional program in patients with urinary tract infections. bioRxiv doi:10.1101/595207:595207.

28. Subashchandrabose S, Hazen TH, Brumbaugh AR, Himpsl SD, Smith SN, Ernst RD, Rasko DA, Mobley HLT. 2014. Host-specific induction of Escherichia coli fitness genes during human urinary tract infection. Proceedings of the National Academy of Sciences 111:18327–18332.

29. Garcia Erin C, Brumbaugh Ariel R, Mobley Harry LT, Payne SM. 2011. Redundancy and Specificity of *Escherichia coli* Iron Acquisition Systems during Urinary Tract Infection. Infection and Immunity 79:1225–1235.

30. Liu X, Matsumura P. 1994. The FlhD/FlhC complex, a transcriptional activator of the *Escherichia coli* flagellar class II operons. J Bacteriol 176:7345–51.

31. Macnab RM. 1996. Flagella and motility. Escherichia coli and Salmonella: cellular and molecular biology:123–145.

32. McCarter LL. 2006. Regulation of flagella. Current Opinion in Microbiology 9:180–186.

33. Martínez-Antonio A, Janga SC, Thieffry D. 2008. Functional organisation of *Escherichia coli* transcriptional regulatory network. Journal of Molecular Biology 381:238–247.

34. Li M, Ho PY, Yao S, Shimizu K. 2006. Effect of lpdA gene knockout on the metabolism in Escherichia coli based on enzyme activities, intracellular metabolite concentrations and metabolic flux analysis by 13C-labeling experiments. Journal of Biotechnology 122:254–266.

35. Quail MA, Haydon DJ, Guest JR. 1994. The *pdhR–aceEF–lpd* operon of *Escherichia coli* expresses the pyruvate dehydrogenase complex. Molecular Microbiology 12:95–104.

36. Hagberg L, Engberg I, Freter R, Lam J, Olling S, Svanborg Edén C. 1983. Ascending, unobstructed urinary tract infection in mice caused by pyelonephritogenic *Escherichia coli* of human origin. Infection and Immunity 40:273.

37. Subashchandrabose S, Mobley HLT. 2015. Back to the metal age: battle for metals at the host-pathogen interface during urinary tract infection. Metallomics : integrated biometal science 7:935–942.

38. Forsyth VS, Himpsl SD, Smith SN, Sarkissian CA, Mike LA, Stocki JA, Sintsova A, Alteri CJ, Mobley HLT. 2020. Optimization of an Experimental Vaccine To Prevent *Escherichia coli* Urinary Tract Infection. mBio 11:e00555–20.

39. Mike LA, Smith SN, Sumner CA, Eaton KA, Mobley HLT. 2016. Siderophore vaccine conjugates protect against uropathogenic *Escherichia coli* urinary tract infection. Proceedings of the National Academy of Sciences 113:13468–13473.

40. Adler J, Templeton B. 1967. The effect of environmental conditions on the motility of *Escherichia coli*. J Gen Microbiol 46:175–84.

41. Guzzo A, Diorio C, DuBow MS. 1991. Transcription of the *Escherichia* coli *fliC* gene is regulated by metal ions. Applied and Environmental Microbiology 57:2255–2259.

42. Hantke K. 1981. Regulation of ferric iron transport in Escherichia coli K12: isolation of a constitutive mutant. Mol Gen Genet 182:288–92.

43. Bagg A, Neilands JB. 1987. Ferric uptake regulation protein acts as a repressor, employing iron (II) as a cofactor to bind the operator of an iron transport operon in Escherichia coli. Biochemistry 26:5471–7.

44. Massé E, Vanderpool CK, Gottesman S. 2005. Effect of RyhB small RNA on global iron use in Escherichia coli. J Bacteriol 187:6962–71.

45. Abdel-Hamid AM, Attwood MM, Guest JR. 2001. Pyruvate oxidase contributes to the aerobic growth efficiency of *Escherichia coli*. Microbiology 147:1483–1498.

46. Alteri CJ, Smith SN, Mobley HLT. 2009. Fitness of Escherichia coli during Urinary Tract Infection Requires Gluconeogenesis and the TCA Cycle. PLOS Pathogens 5:e1000448.

47. Alteri CJ, Himpsl SD, Mobley HLT. 2015. Preferential Use of Central Metabolism In Vivo Reveals a Nutritional Basis for Polymicrobial Infection. PLOS Pathogens 11:e1004601.

48. Datsenko KA, Wanner BL. 2000. One-step inactivation of chromosomal genes in *Escherichia coli* K-12 using PCR products. Proceedings of the National Academy of Sciences 97:6640.

49. Shea Allyson E, Stocki Jolie A, Himpsl Stephanie D, Smith Sara N, Mobley Harry LT. Loss of an Intimin-Like Protein Encoded on a Uropathogenic *E. coli* Pathogenicity Island Reduces Inflammation and Affects Interactions with the Urothelium. Infection and Immunity 0:IAI.00275–21.

